## Appendix for "SNARE mimicry by the CD225 domain of IFITM3 enables regulation of homotypic late endosome fusion"

Rahman et al. *The EMBO Journal,* 2024

### Appendix

Table of Contents:

Appendix Figure S1……………………………….page 2
Appendix Figure S2……………………………….page 3
Appendix Figure S3……………………………….page 4
Appendix Figure S4……………………………….pages 5-6

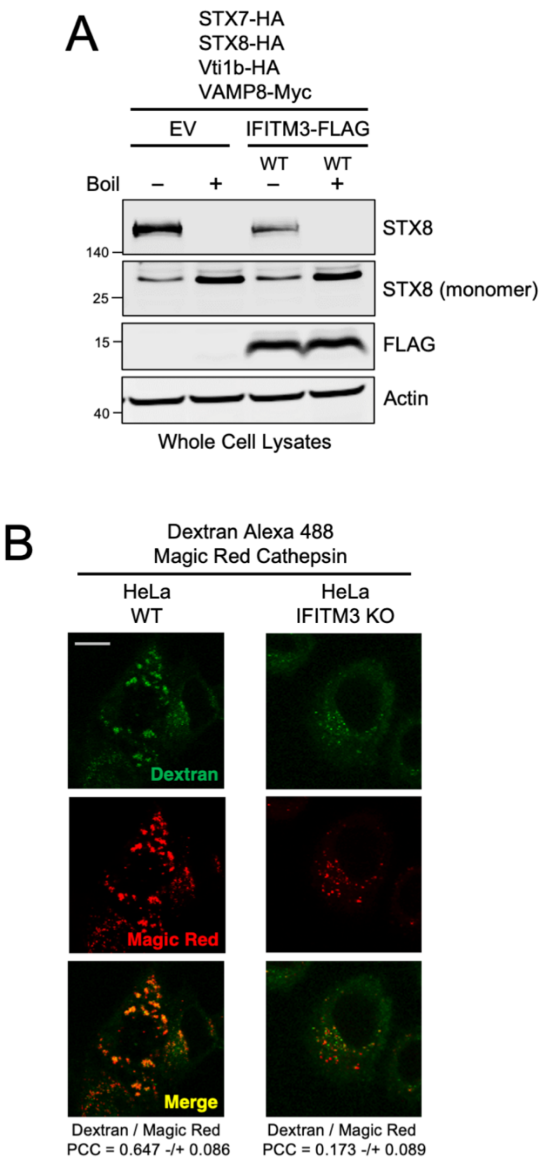


**Appendix Figure S1:**
(A) HEK293T stably expressing Empty Vector, IFITM3 WT-FLAG, or IFITM3 G95L-FLAG were co-transfected with STX7-HA, STX8-HA, Vti1b-HA, and VAMP8-Myc. Following whole cell lysis, samples were either boiled at 100°C (+) or not (-). SDS-PAGE and immunoblotting was performed with anti-STX8 and anti-FLAG. Actin was used as loading control. Numbers and tick marks left of blots indicate position and size (in kilodaltons) of protein standard in ladder. Immunoblots were performed independently twice, and a representative example is shown. (B) HeLa cells (WT or *IFITM3* KO) were pulsed with Dextran Alexa Fluor 488 for 2 hrs followed by addition of Magic Red for 5 mins. Living cells were then analyzed immediately by confocal immunofluorescence microscopy. Colocalization between Dextran and Magic Red was measured by calculating the Pearson's Correlation Coefficient using Fiji software. Coefficients were calculated from medial Z-slices from three fields of view containing 5-15 cells per condition and presented as means and standard error. Scale bar = 15 microns. WT; wild-type. PCC; Pearson's Correlation Coefficient.

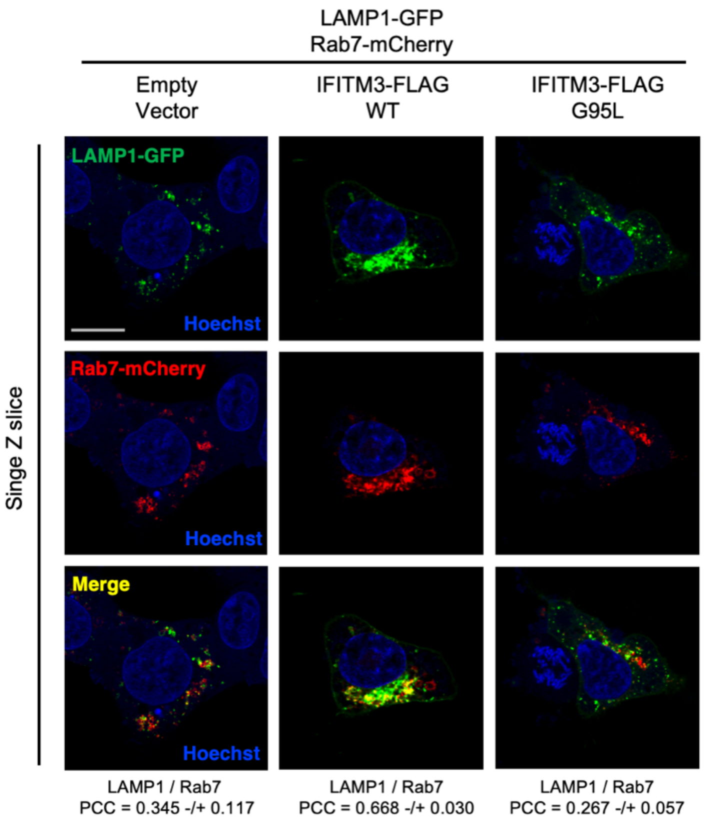


**Appendix Figure S2:**
HEK293T cells stably expressing Empty Vector, IFITM3 WT-FLAG, or IFITM3 G95L-FLAG were transfected with LAMP1-GFP and Rab7-mCherry, fixed, and analyzed by confocal fluorescence microscopy. Hoechst was used to stain nuclei. Colocalization was measured between LAMP1-GFP and Rab7-mCherry by calculating the Pearson's Correlation Coefficient using Fiji software. Coefficients were calculated from medial Z-slices from three fields of view containing 5-15 cells per condition and presented as means and standard error. Scale bar = 15 microns.

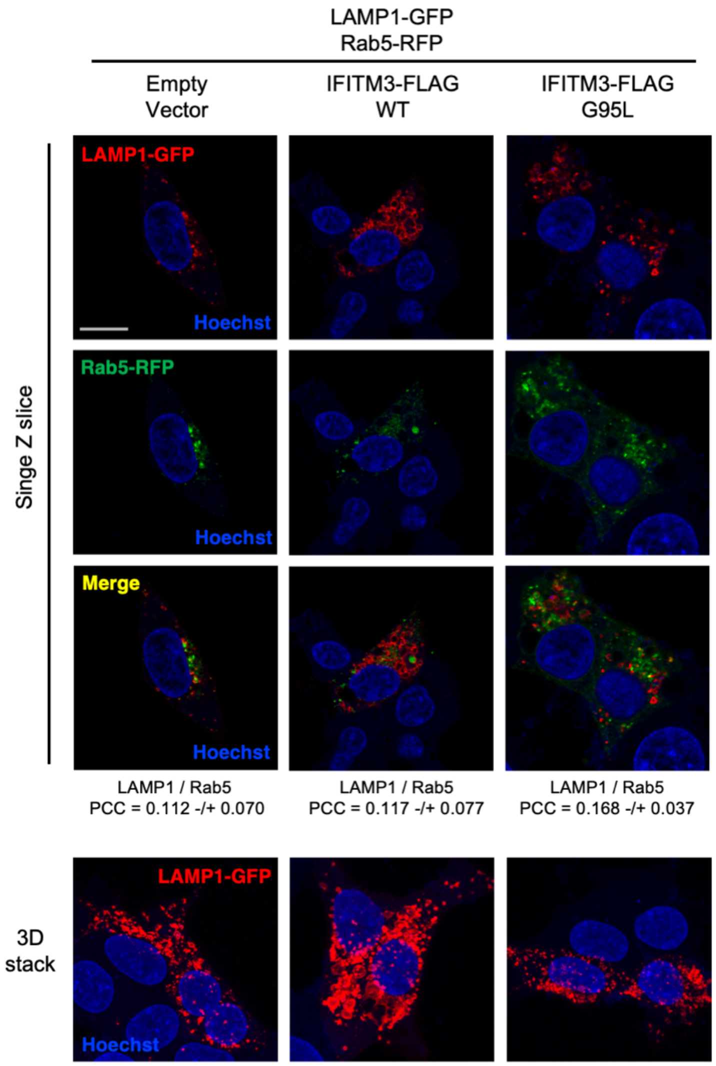


**Appendix Figure S3:**
HEK293T cells stably expressing Empty Vector, IFITM3 WT-FLAG, or IFITM3 G95L-FLAG were transfected with LAMP1-GFP and Rab5-RFP, fixed, and analyzed by confocal fluorescence microscopy. Hoechst was used to stain nuclei. Colocalization was measured between LAMP1-GFP and Rab5-RFP by calculating the Pearson's Correlation Coefficient using Fiji software. Coefficients were calculated from medial Z-slices from three fields of view containing 5-15 cells per condition and presented as means and standard error. Scale bar = 15 microns.

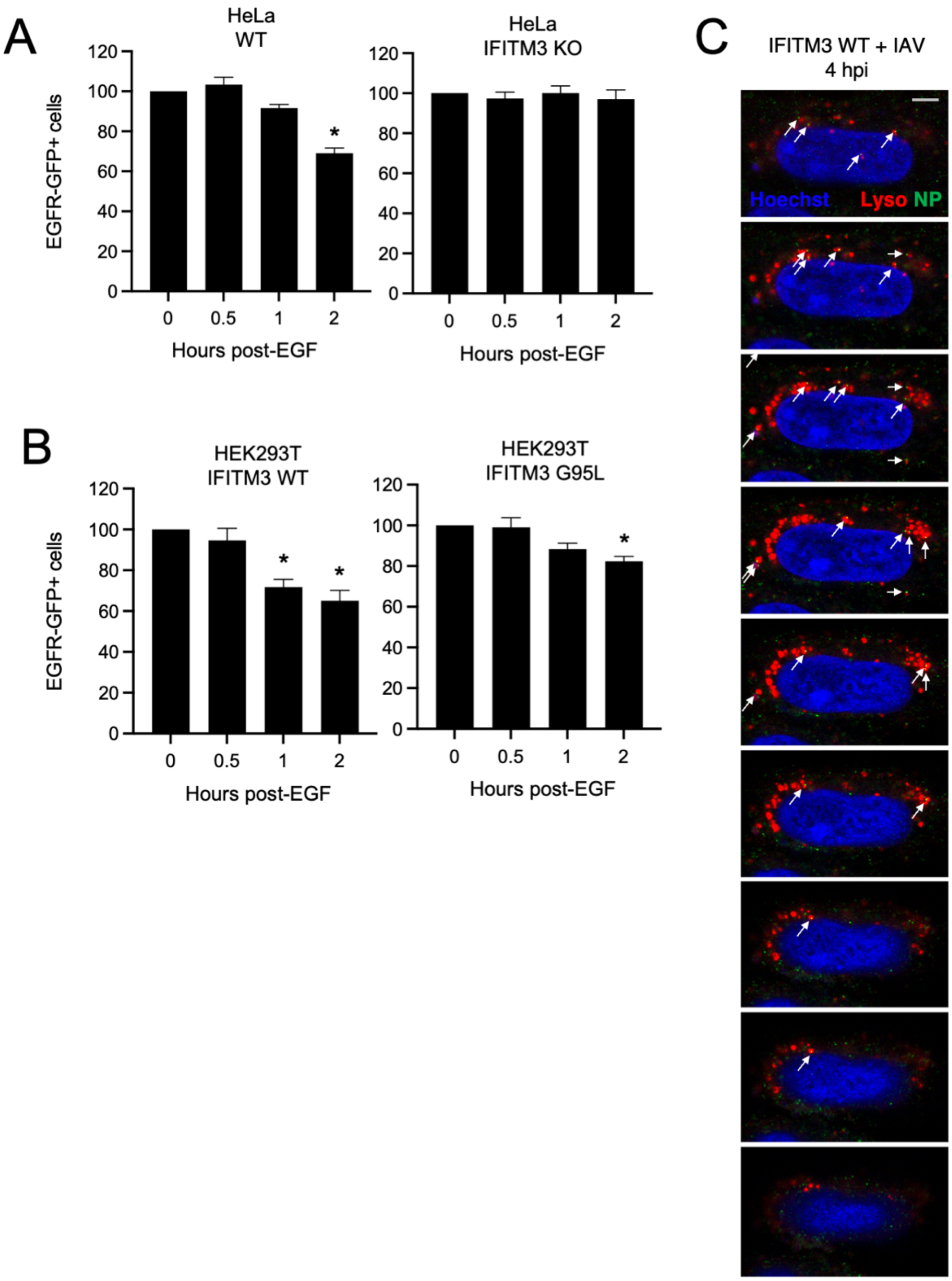


**Appendix Figure S4:**
(A) HeLa cells (WT or *IFITM3* KO) were transfected with EGFR-GFP and treated with EGF. At the indicated time points, EGFR-GFP+ cells were analyzed by flow cytometry. The percentage of EGFR-GFP+ cells was normalized to 100% at time 0 post-EGF addition. Statistically significant differences between the indicated condition at time 0 were determined by one-way ANOVA (*, $p < 0.0001$) (B) HEK293T cells stably expressing IFITM3 WT or IFITM3 G95L were transfected with EGFR-GFP and treated with EGF. At the indicated time points, EGFR-GFP+ cells were analyzed by flow cytometry. The percentage of EGFR-GFP+ cells was normalized to 100% at time 0 post-EGF addition. Statistically significant differences between the indicated condition and time

0 were determined by one-way ANOVA (*, $p < 0.01$). (C) An additional example of IAV trafficking to lysosomes in cells expressing IFITM3 WT. A succession of Z slices from the confocal stack are shown. White arrows indicate bright NP puncta that colocalize with Lysotracker or which tightly appose a Lysotracker+ compartment. Scale bar = 5 microns. WT; wild-type. hpi; hours post-infection. Lyso; Lysotracker.
