## Supplementary figures and images for "SNARE mimicry by the CD225 domain of IFITM3 enables regulation of homotypic late endosome fusion"

### Figure EV1

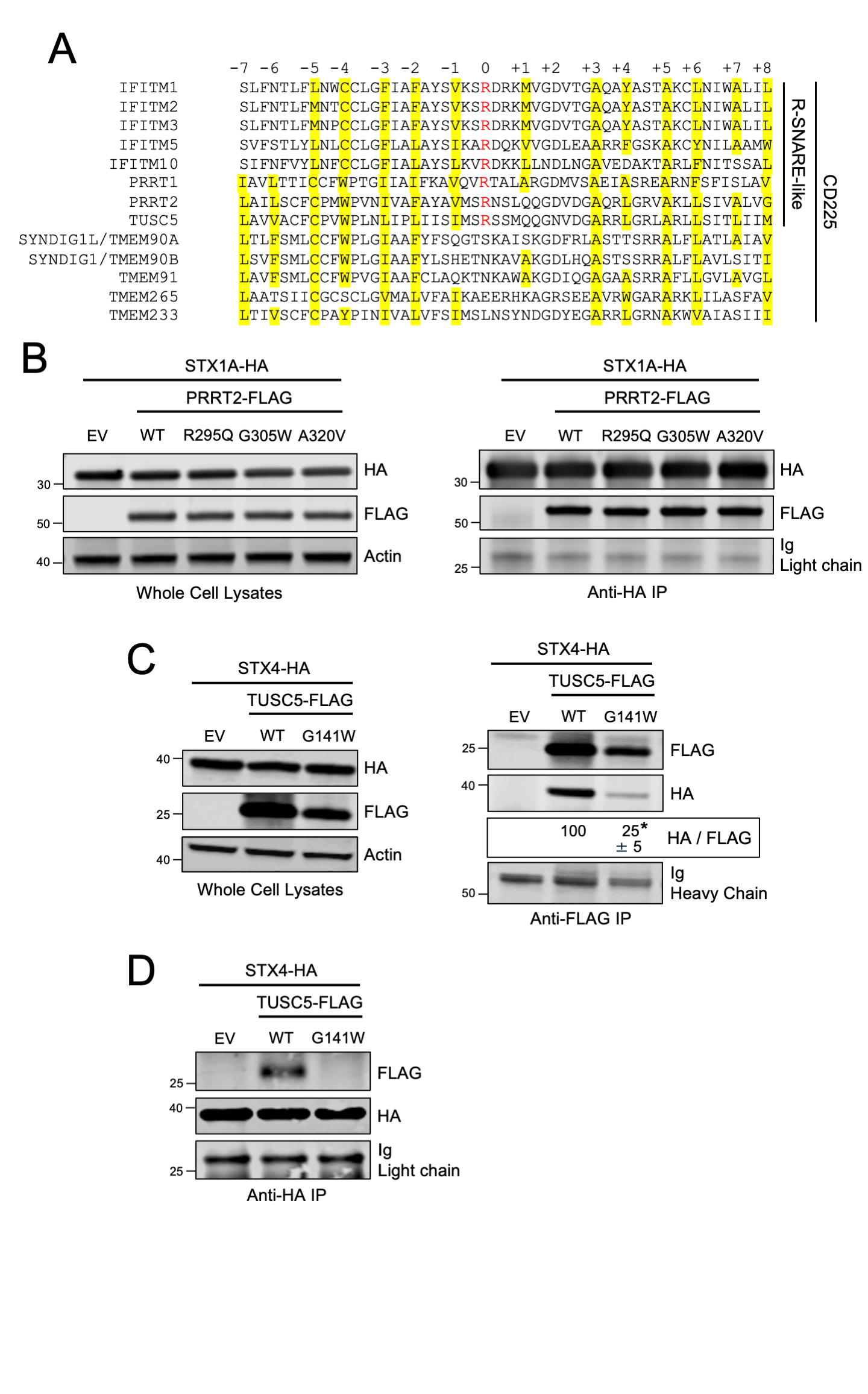

### Figure EV2

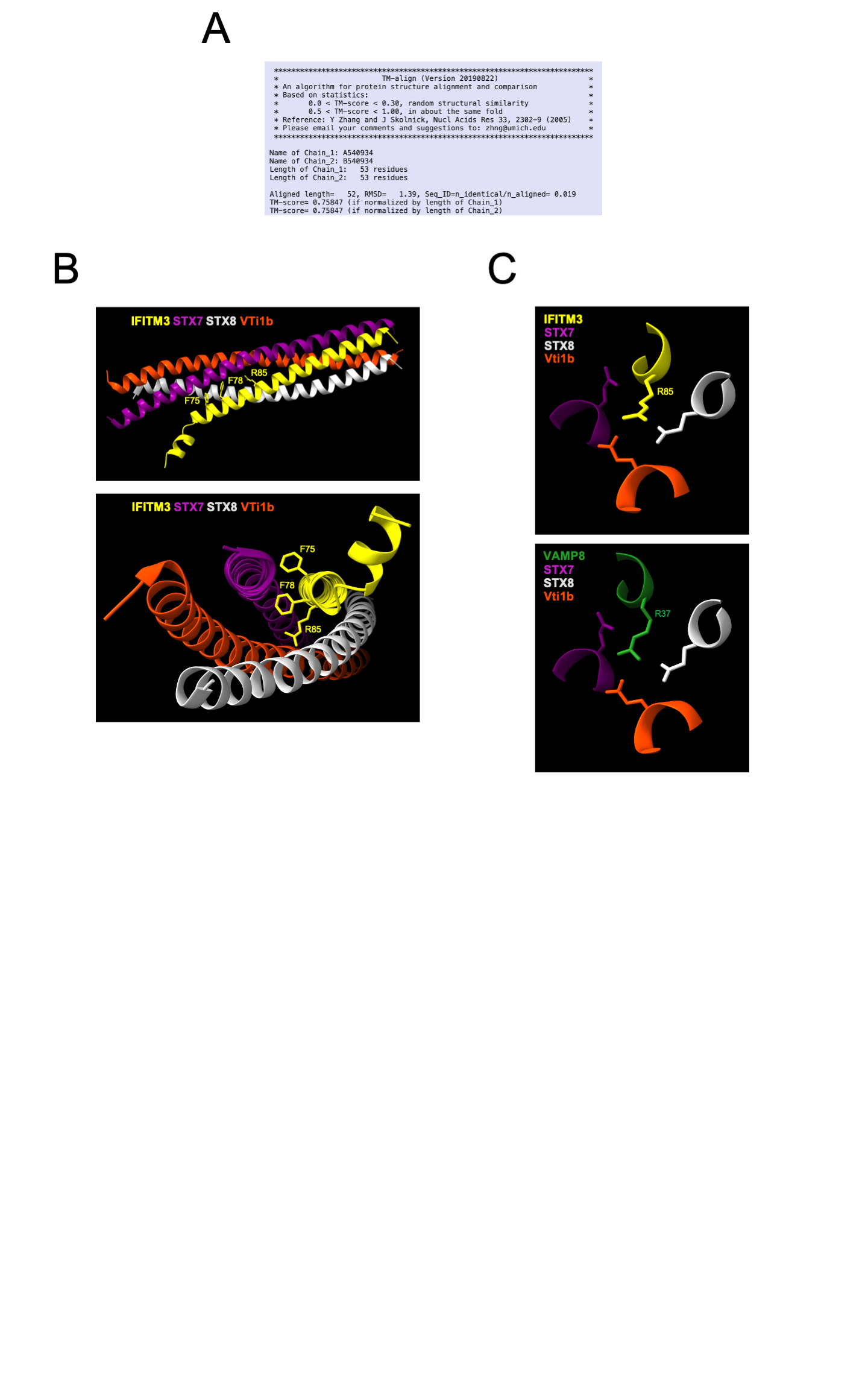

### Figure EV3

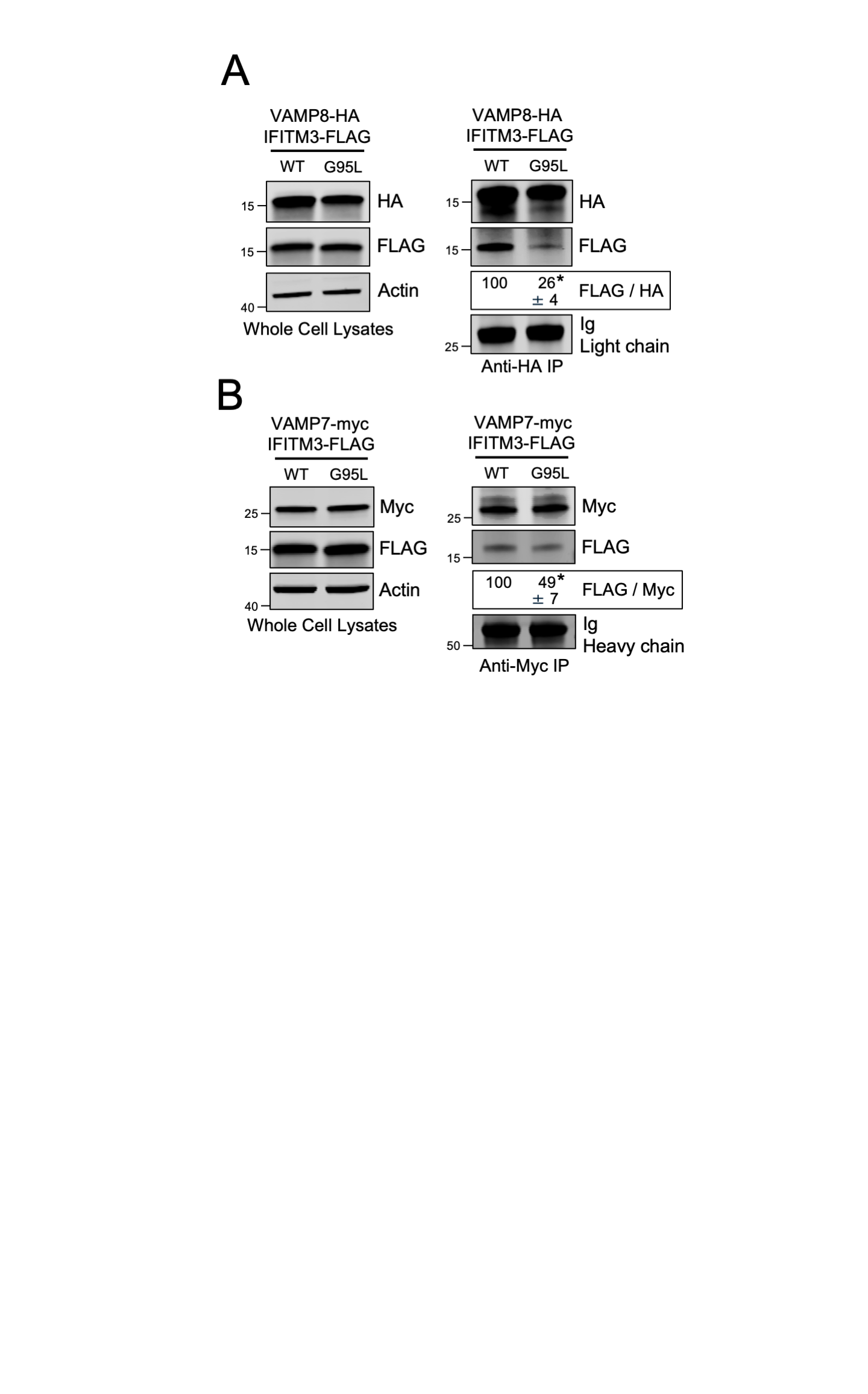

### Figure EV4

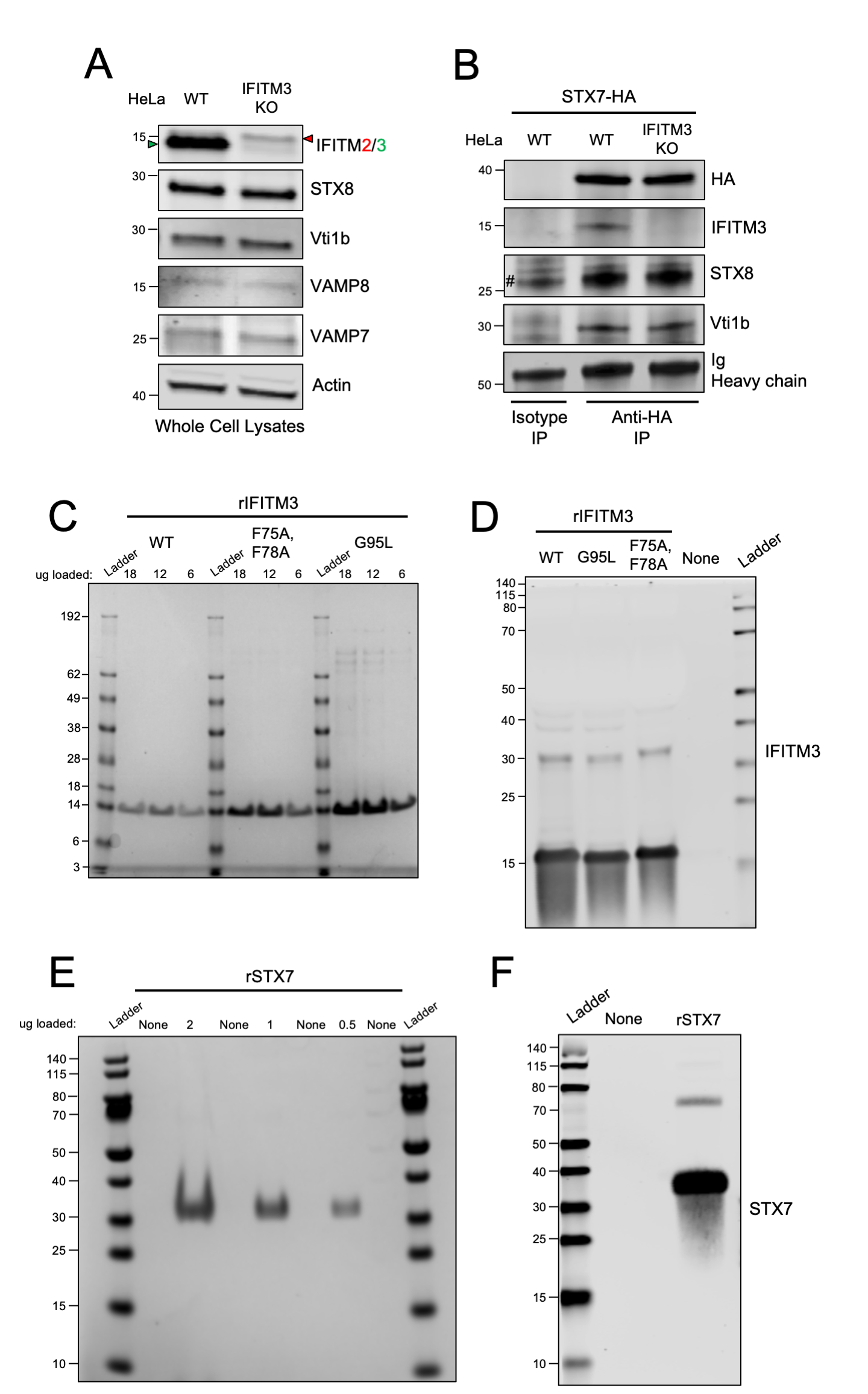

### Figure EV5

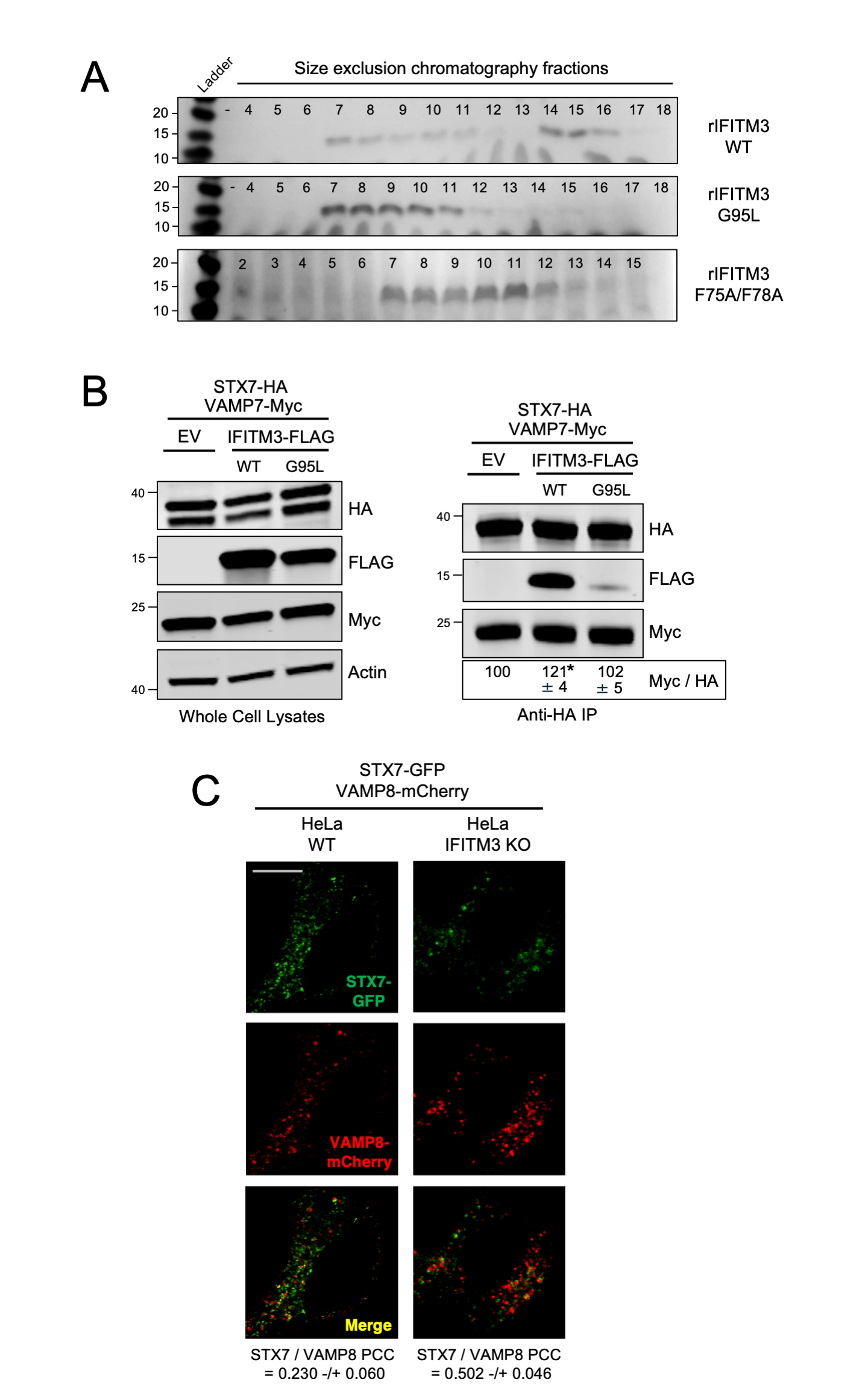
